# Sensitivity to the slope of the amplitude spectrum is dependent on the spectral slopes of recently viewed environments: A visual adaptation study in modified reality

**DOI:** 10.1101/2021.08.19.456985

**Authors:** Bruno Richard, Patrick Shafto

**Affiliations:** Department of Mathematics and Computer Science, Rutgers University - Newark, 101 Warren Street, Rm 216, Newark, New Jersey, USA, 07102; School of Mathematics, Institute for Advanced Study, Princeton, NJ

**Keywords:** Natural Scenes, Amplitude Spectrum Slope, Head-Mounted Display, Modified Environment, Bayesian Observer

## Abstract

Scenes contain many statistical regularities that could benefit visual processing if accounted for by the visual system. One such statistic is the orientation-averaged slope (*α*) of the amplitude spectrum of natural scenes. Human observers show different discrimination sensitivity to *α*: sensitivity is highest for *α* values between 1.0 and 1.2 and decreases as *α* is steepened or shallowed. The range of *α* for peak discrimination sensitivity is concordant with the average *α* of natural scenes, which may indicate that visual mechanisms are optimized to process information at *α* values commonly encountered in the environment. Here we explore the association between peak discrimination sensitivity and the most viewed *α*s in natural environments. Specifically, we verified whether discrimination sensitivity depends on the recently viewed environments. Observers were immersed, using a Head-Mounted Display, in an environment that was either unaltered or had its average *α* steepened or shallowed by 0.4. Discrimination thresholds were affected by the average shift in *α*, but this effect was most prominent following adaptation to a shallowed environment. We modeled these data with a Bayesian observer and explored whether a change in the prior or a change in the likelihood best explained the psychophysical effects. Change in discrimination thresholds following adaptation could be explained by a shift in the central tendency of the prior concordant with the shift of the environment, in addition to a change in the likelihood. Our findings suggest that expectations on the occurrence of *α* that result from a lifetime of exposure remain plastic and able to accommodate for the statistical structure of recently viewed environments.

---

Our visual world is diverse but contains spatial regularities that span across the various environments we encounter. There is evidence that accounting for these spatial regularities in encoding can improve the visual system’s efficiency in processing complex scene properties [1, 2, 3, 4]. One such regularity is the slope of the amplitude spectrum (*α*) of natural scenes, which defines the association of amplitude to spatial frequency (*f*),

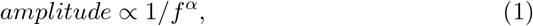

where the average *α* value across various natural scenes is 1.08 [5, 6, 7, 8, 9, 10, 11]. Observers are sensitive to and influenced by the value of *α*: they exhibit different discrimination and recognition abilities for different *α* values [1, 12, 13, 14] and can adapt to *α* [15, 16, 17]. This bias in the discrimination ability of observers for *α* may be associated with the recent visual experience of observers [15]. Here, we explore how the discrimination sensitivity for *α* is related to the distribution of *α*s gathered from recent perceptual experience. We used a Head-Mounted Display (HMD) system paired with a camera to display and modify the natural visual environment of observers and measure discrimination thresholds to *α*. Finally, we interpret our findings according to a Bayesian observer model [18, 19, 20] and derive the shape and malleability of the prior for the αs of natural scenes.

Measuring the discriminability of *α* has been a popular method for determining the shape of the internal representation for *α* [9, 21, 22, 23]. For example, *α* common finding when measured with noise stimuli – modified to have a particular *α* value, – is that the discriminability of *α* is best (thresholds are lowest) when the reference *α* is between 1.0 and 1.3, and rises as the *α* is steepened or shallowed [24, 13, 25, 13], a range of *α* near the average *α* of natural scenes. Thus it appears that the human visual system is best able to discriminate the spatial characteristics of scenes it encounters often. However, the peak in discriminability can depend on the original *α* value of the image. When observers are asked to discriminate against the original *α* of a natural image, thresholds are significantly higher than when observers complete the discrimination task against an image with a modified *α*^1^ [26]. Tolerance for small deviations in the natural *α* of an image [26, 22, 23] may assist in optimizing visual processing by facilitating the discrimination of various objects in the natural world [12, 14]. Thus, while the human visual system peak sensitivity to *α* is for values regularly encountered (*α* ≈ 1.0), it is simultaneously tolerant for minor deviations from the natural *α* of an image, which suggests some process of adaptation to the spatial characteristics of the environment.

Visual adaptation is the process by which exposure to a stimulus affects visual processing for subsequent stimuli [27, 28, 29, 30]. When the adapting stimuli are spatially complex, like a noise stimulus modified to have a natural *α* value, adaptation acts to renormalize the percept of observers towards an internal norm; a neutral point in the responses of the detecting mechanisms [15]. Adaptation to artificially steepened or shallowed images generates a shift in the perceived *α* of a new image in the opposite direction to that of the adapting stimulus *α*. No shift in the perceived *α* of images is measured when the adapting stimulus has an *α* of 1.0, which is coincidentally the average value of *α* across all natural scenes [15]. A finding that indicates the human visual system has developed an internal representation or an expectation for *α* values commonly encountered through visual experience.

The statistical structure of natural scenes is known to influence sensitivity to visual features [31, 19, 20, 32]. For example, there is an orientation anisotropy in human vision whereby sensitivity to horizontal content embedded in natural scenes is worse than that of obliques (i.e., the horizontal effect; [33, 34, 35]). The horizontal effect is believed to stem from the over-representation of horizontal content in natural scenes that is perceptually normalized by reducing sensitivity to horizontal content [33, 11, 19, 34]. If observers are immersed in an environment that does not contain the natural distribution of orientation content, the strength of the horizontal effect diminishes severely [31]. This reduction in the magnitude of the horizontal effect occurs relatively rapidly (within an hour or two of adaptation), which is a strong indication that the expected structure of natural scenes can be relearned with a relatively brief exposure to a novel environment [32, 31]. The change in sensitivity to orientation contrast was well-explained by changing the prior of a Bayesian observer to match the novel orientation distribution of the adapting environment [31]. If the internal representation for *α* is determined by the distribution of commonly viewed *α*s in the natural environment, then it may be able to adjust given a novel environment statistical structure, as has been demonstrated with orientation sensitivity in natural scenes.

Here, we explore whether sensitivity to *α* is associated with the statistical structure, or distribution, of *α* in natural scenes by measuring *α* discrimination thresholds prior to and following immersion in an environment that no longer contains the typical *α* distribution of scenes. Observers were immersed in the environment with a Head-Mounted Displays (HMDs) and a camera that recorded and displayed the visual world in near real-time (a display system that we have named Modified Reality; [31]). All experimental conditions, the measurement of *α* discrimination thresholds and the adaptation to the modified environment were conducted in the Head-Mounted Display. This method greatly facilitates the measurement of sensitivity to features in the natural environment of observers while maintaining sufficient experimental control on the presented stimuli. The environment was modified by steepening or shallowing the distribution of *α*s. Finally we develop a Bayesian observer model to explore the possible shape of a prior for *α* and verify if the prior adjusts to match the novel environmental distribution of *α* [31, 19, 18].

## Materials and Methods

### Ethics Statement

Procedures were approved by the Rutgers Arts and Sciences Institutional Review Board (Study ID: Pro2020002981). The experiment was completed by three male participants who provided written informed consent.

### Participants

Three male observers (age 29, 33, and 34) took part in this study including one author (BR). All participants were experienced psychophysical observers with prior training on the amplitude spectrum slope discrimination task and had normal or corrected to normal visual acuity.

### Apparatus

The experiment was conducted in an *Oculus Rift Development Kit 2* (DK2). The DK2 has two 5.7 inch OLED displays, each having a monocular resolution of 960×1080 pixels and a refresh rate of 75Hz. The Field of View of the Oculus Rift DK2 is 100°, resulting in a visual resolution of approximately 11 pixels per degree of visual angle^2^. The observer’s environment was captured by a *See3Cam_130* USB 3.0 camera mounted on the Oculus Rift. The camera continuously collected images of the observers’ environment at a rate of 60 fps during the experiment. The images recorded by the camera were sent to the laptop computer (Dell Latitude 3470), using the MATLAB Image Acquisition Toolbox, filtered in MATLAB and displayed back to the Oculus Rift with the the Oculus VR library of *Psychtoolbox* [36, 37].

### Stimuli

The environment images collected by the mounted camera were converted to grayscale and cropped to a size of 720×1080 pixels. The 120 pixels of the Oculus display above and below the presented environment were set to mean luminance (RGB [128,128,128]). At baseline condition, only the RMS contrast of the environment images was adjusted to match that of the target stimuli (RMS = 0.15). For the adaptation condition, images collected from the environment had their global *α* changed by either +0.4 or −0.4 (see **Figure 1**). The environment *α* was changed in an identical manner to the creation of the synthetic noise images used to measure discrimination thresholds (**Figure 1**). The amplitude of each spatial frequency coefficient was divided by the average amplitude (calculated across orientation for that spatial frequency) to create a flat (α = 0.0) amplitude spectrum [22]. In this form, the *α* of the environment image was adjusted by multiplying each spatial frequency’s coefficient by *f^-α_adjust_^* (the original image *α* + 0.4 for the steeper condition or – 0.4 for the shallower condition). The RMS contrast of *α_adjust_* images was then adjusted to 0.15.

**Figure 1:**
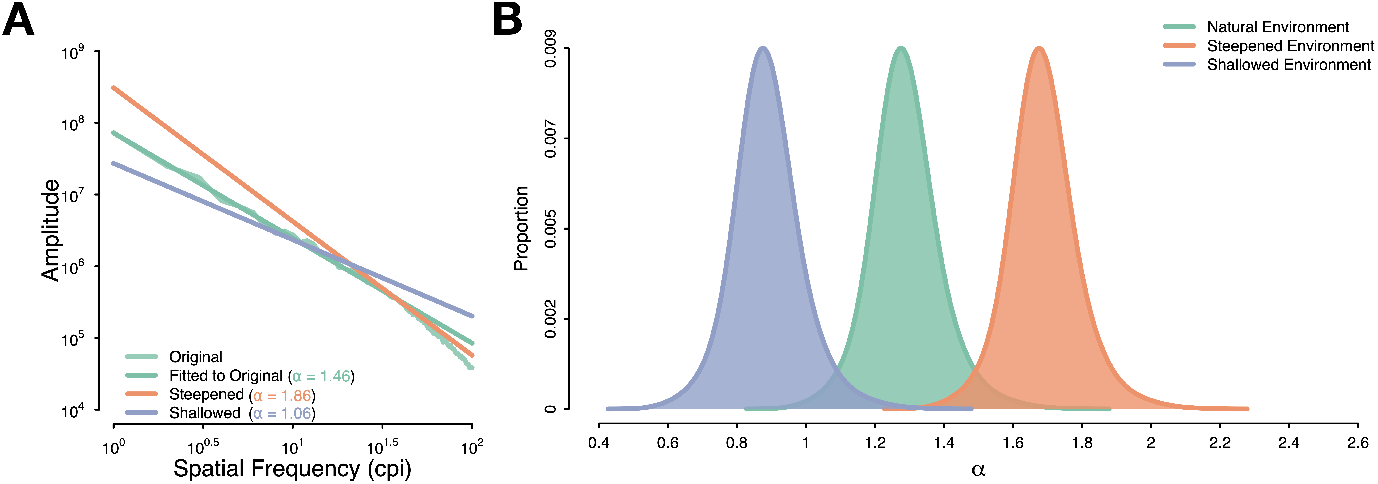
**A** *α* modification procedures: We fit a linear regression to the log orientation averaged amplitude spectrum of the original image (Original) to measure its *α_original_*. The new amplitude spectrum slope was created by first dividing the spatial frequency coefficients by the average amplitude to create a flat amplitude spectrum. The *α* of the original image could then be adjusted by multiplying each spatial frequency coefficient by *α_original_* + 0.4. The modified spectrum was then scaled vertically to ensure that the total contrast energy in the modified image was identical to that of the original. **B** The distribution of *α* values in the environment of our observers. The α distribution of our laboratory was similar to that reported by [6] for fully carpented environments with a mean *α* = 1.29 and standard deviation of 0.09. The steep environment distribution has a mean of 1.69 while the shallow environment had a mean of 0.89.

All test stimuli consisted of synthetic visual noise patterns constructed in the Fourier domain using MATLAB (Mathworks, Natick, MA) and corresponding Signal Processing and Image Processing toolboxes. We opted for noise patterns with different *α* values as test stimuli, and not natural images, as these were overlaid onto the modified environment when we measured discrimination thresholds and this greatly simplifies the identification of the test stimuli for observers. The visual noise stimuli were created by constructing a polar matrix for the amplitude spectrum and assigning all coordinates the same arbitrary amplitude coefficient (except at the location of the DC component, which was assigned a value of 0). The result is a flat isotropic broadband spectrum (i.e., *α* = 0.0), referred to as the template amplitude spectrum [24, 22]. In this form, the *α* of the template spectrum can be adjusted by multiplying each spatial frequency’s amplitude coefficient by f^-*α*^. The phase spectra of test stimuli were constructed by assigning random values from -*π* to *π* to the different coordinates of a polar matrix while maintaining an odd-symmetric phase relationship to maintain conjugate symmetry. The noise patterns were rendered into the spatial domain by taking the inverse Fourier transform of an α-altered template amplitude spectrum and a given random phase spectrum. The phase spectrum for all stimuli presented within a trial was identical but randomized from trial-to-trial. The 1/f^*α*^ stimuli were 20° in size (220 pixels). RMS contrast (the standard deviation of all pixel luminance values divided by the mean of all pixel luminance values) was fixed to 0.15.

### Procedures

The experimental procedures measured sensitivity to *α* prior to and following adaptation to a modified environment. Participants first completed an *α* discrimination task with 1/f*^α^* noise stimuli placed atop the unaltered environment (RMS adjusted only; see **Figure 2)**. Discrimination thresholds were estimated by a temporal three-interval, two-alternative “Odd-Man-Out” forced choice task [24, 13, 25]. The trial-to-trial change in the image’s *α* was controlled by a 1- up, 2-down staircase procedure targeting 70.71% correct [38, 39]. The staircase began with a difference between the odd and reference stimulus of 0.5, which decreased in linear steps (step size = 0.02) towards the reference *α* when the observer made two consecutive correct responses. The difference was increased by 0.02 when observers made an incorrect response (1-Up / 2-Down Rule).

**Figure 2:**
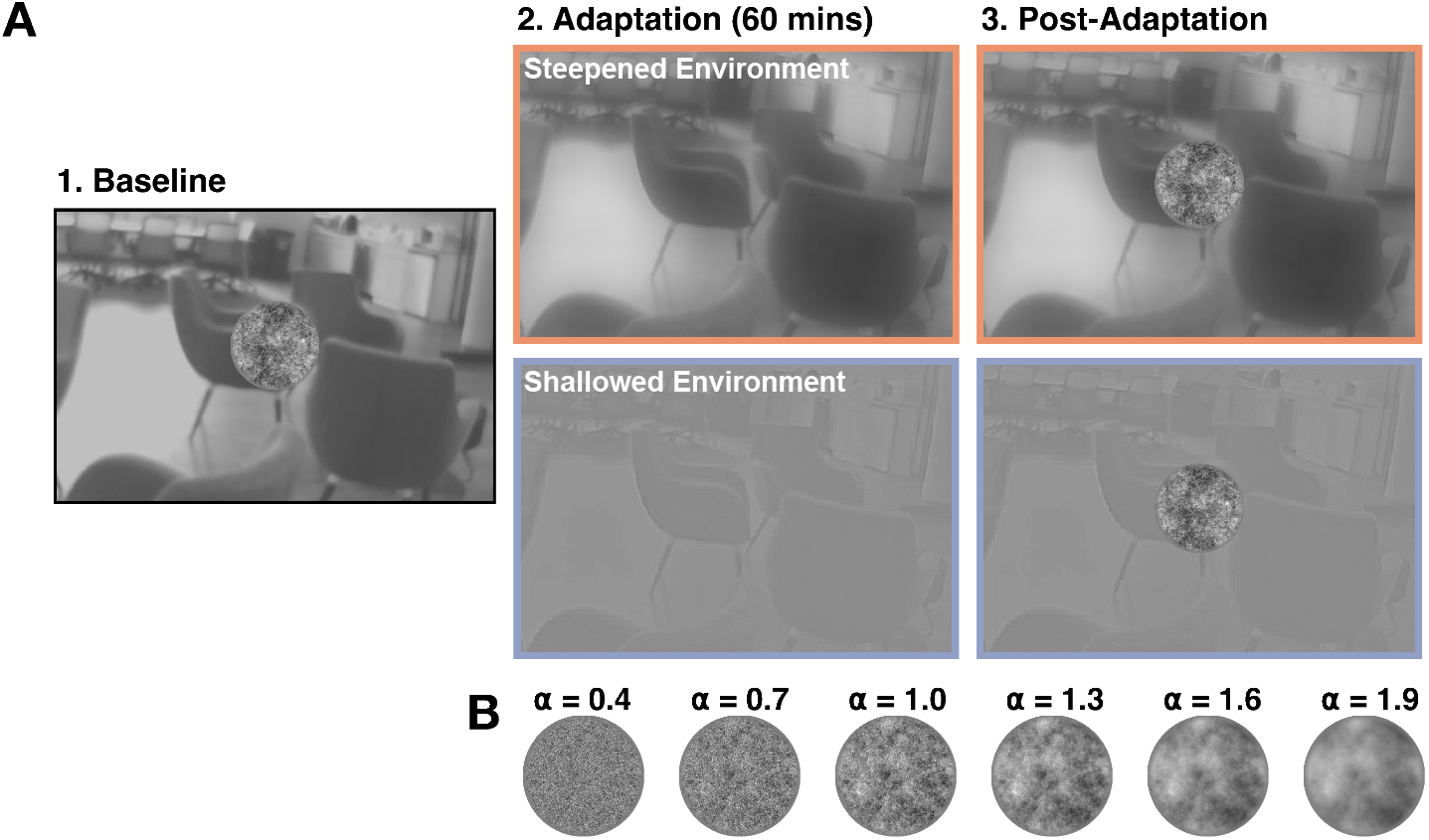
**A** Prior to adaptation, participants completed a baseline *α* discrimination task in their unaltered (gray-scaled and RMS contrast adjusted to 0.15). Participants did 2 staircases per reference *α* [0.4, 0.7, 1.0, 1.3, 1.6] (see **B** for example reference *α*). Participants then began the adaptation procedure, where they viewed the modified world (global *α* of the environment were steepened by +0.4) for a period of 60 minutes. Following adaptation, participants completed the same *α* discrimination task as baseline, but with the modified environment in the background. Note that the contrast of the test pattern has been increased for visibility.

Each trial began with a black (RGB [0, 0, 0]) fixation point (0.5°) presented for 1000ms at the center of the display, followed by three stimulus presentation intervals (250ms each) interlaced by a blank screen (environment only) for 500ms. The second interval always contained the reference *α* while either the first or third contained the odd stimulus (the other being identical to the reference *α* interval). Participants indicated which interval, either the first or the third, they perceived as being the “Odd-Man-Out” via keyboard press. The staircase continued until 12 reversals occurred and thresholds were estimated by averaging the *α* values of the odd stimulus for the last 5 reversals. Observers completed two staircases per reference *α* value [0.4, 0.7, 1.0, 1.3, 1.6, 1.9]; the discrimination thresholds of each were averaged for data analysis.

Immediately following baseline, participants completed a 60 minute adaptation period immersed in the modified environment (either *α* shifted by +0.4 or −0.4) in the HMD. Participants were instructed to look around their environment and interact with objects and individuals present around them but to remain seated or standing in place. Observers then repeated the *α* discrimination task with the modified environment presented in the background. Participants repeated the experiment with the other modified environment (either *α* shifted by +0.4 or −0.4) no less than 24 hours following the first experimental session. Each experimental session took approximately 2.5 hours to complete (45 minutes per *α* discrimination task and 60 minutes of adaptation).

## Results

Average *α* discrimination thresholds measured at baseline, following the adaptation conditions (+0.4 and −0.4), and resulting model fits are shown in **Figure 3**. Discrimination thresholds measured in the Oculus Rift DK2 follow the typical peak and trough commonly observed with traditional displays (i.e., CRTs) [24, 13, 25, 22, 21]. While the overall effects of adaptation on discrimination thresholds are small, we observe some meaningful changes in discrimination according to the adaptation condition. Adaptation to a steepened environment lead to a reduction in discrimination thresholds to very steep reference *α* (1.9), *t*(2) = −3.48, *p* = .037, g = −0.57 [-1.17, 0.05]. Adaptation to a shallowed environment reduced discrimination thresholds for a reference *α* of 0.7, and while this decrease was very large, it was not statistically significant, t(2) = −1.84, *p* = .104, g = −1.58 [−3.08, 0.72]. Given the small number of participants in this study, it is difficult to interpret our findings using traditional null hypothesis significance testing and thus we opt for a descriptive approach with effect size measures (Hedge’s g) and their exact 95% confidence intervals [40], with values reported in **Table 1.** We find steep adaptation to generate a small decrease in discrimination thresholds for reference *α* values below 1.3 and a moderate decrease in discrimination thresholds at steep reference *α* values (*α* = 1.9). Shallow adaptation effects were concentrated at shallow reference *α*s, with a moderate decrease in discrimination thresholds values of 0.4 and 0.7. While the adaptation effects are moderate in magnitude, they nevertheless suggest some influence of environmental *α* on human sensitivity to *α*, whereby sensitivity improved for α values near those of the adapter [41, 15, 42, 43].

**Figure 3:**
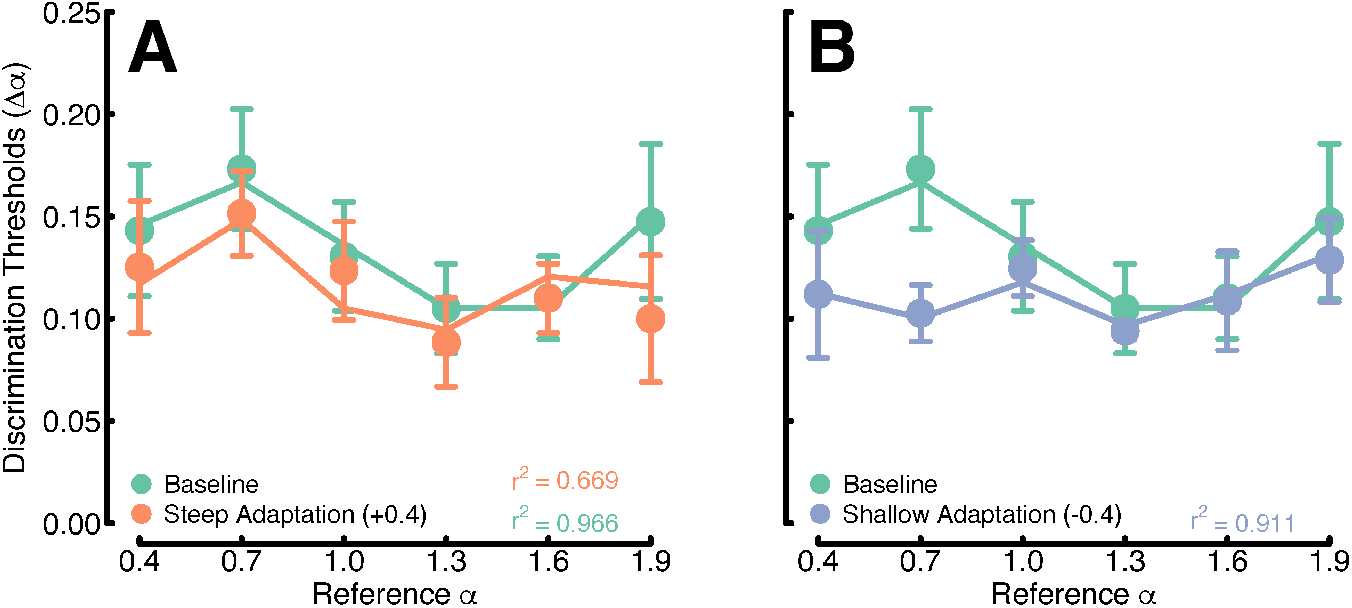
Average discrimination thresholds (points) and environmental priors model fit (lines) for the six reference *α*s used in this study. **A**. Discrimination thresholds measured at baseline (green) and following steep adaptation (orange). **B**. Discrimination thresholds measured at baseline (green) and following shallow adaptation (blue).Adaptation conditions are drawn on separate charts for clarity. Error bars represent ± 1 standard error of the mean. The environmental priors model captured observer discrimination thresholds very well at baseline (*r*^2^ = 0.966) and following adaptation to a shallow environment (*r*^2^ = 0.911) while it struggled with the steep adaptation condition (*r*^2^ = 0.669).

**Table 1:**
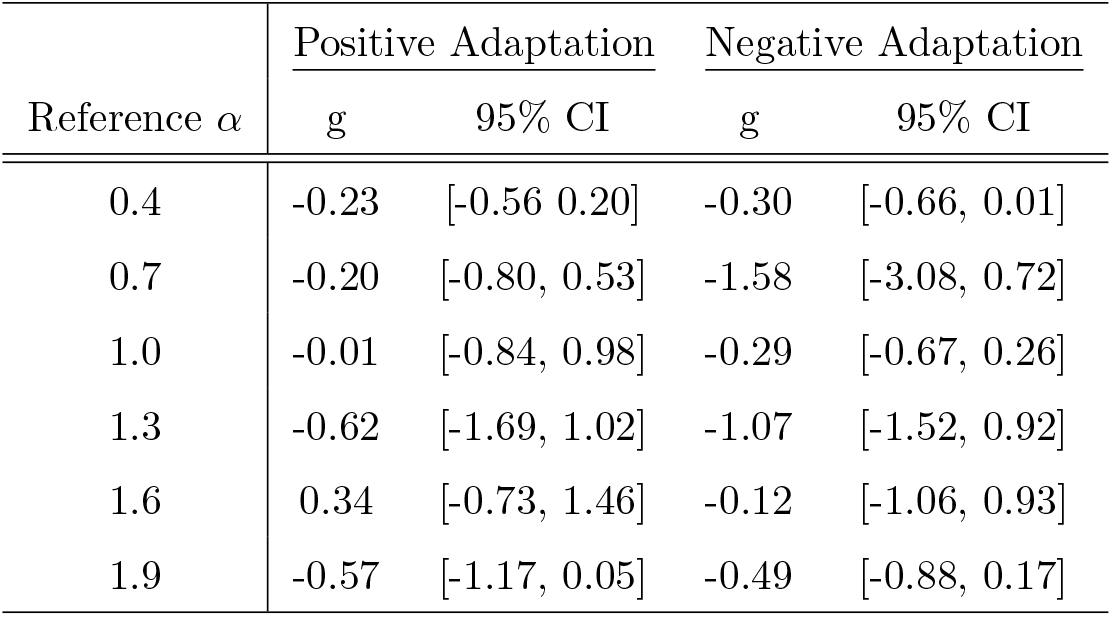
The effect size (Hedge’s g) and exact 95% confidence intervals [40] for differences in the average discrimination following either the positive (+0.4) or negative (−0.4) adaptation conditions.

In both adaptation conditions, we find a facilitation in discrimination towards the average *α* of the adapting environment. Under a Bayesian observer framework, this change in discrimination thresholds could be attributed to an improvement in measurement reliability: a change in the likelihood [42, 44, 43] or in the prior [31] to match the novel environment distribution. In this context, adaption increases the signal-to-noise ratio near the adapter – a change in the likelihood of the Bayesian observer – which increases sensitivity to similar stimulus values. Our results could also be attributed to a change in expectation on the occurrence of *α* values following adaptation – a change in the prior of the Bayesian observer. A change in the prior for *α* may not be far-fetched. An interesting factor to consider is that the mode of *α* distributions varies according to the type of environment they represent (e.g., human-made versus natural) [6]. Different *α* values will thus vary in occurrence when observers are in entirely man-made, natural, or hybrid environments. Previous work has shown that the shape of the prior for statistical regularities in natural scenes (i.e., orientation contrast) matches the distribution of the statistical regularity [19] and if the distribution is changed [31], a similar change in the prior can be observed. We built a Bayesian observer model to determine whether a change in experience (i.e., prior) or an improvement in measurement (i.e., likelihood) accounts for the change in discrimination thresholds we observed here.

The probability of observing a particular stimulus, *α*, given a measurement, *m*, is defined as the product of the likelihood, *p*(*m|α*), and prior probability distribution *p*(*α*)

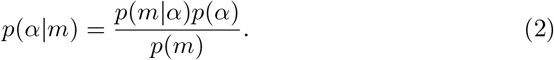

We explored five priors probability distributions to determine if a change in the prior or a change in the likelihood best captured the observed change in discrimination thresholds. The priors were all defined as Gaussian distributions,

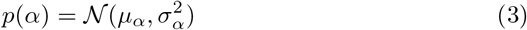

with their means *μ_α_* set to one of the following conditions. In the first condition, the mean value of the priors matched those of the adaptation environment (the envrionmental priors model; baseline: *μ_α_* = 1.29, steep adaptation: *μ_α_* = 1.69, shallow adaptation: *μ_α_* = 0.89), the second to the fourth alternatives were models with a prior fixed at one of the environmental distributions, (baseline, steep or shallow models) while the fifth alternative had a prior mean that matched that the distribution of *α*s recorded from a variety of environments (the natural environment prior; *α* = 1.08). The standard deviation (*σ_α_*) of all priors was fixed to 0.15, slightly broader than the recorded environmental prior but a better approximation of the width of *α* distributions recorded in various environments [5].

The observer likelihood *p*(*m|α_i_*) was defined as a Gaussian centered on the current *α_i_*,

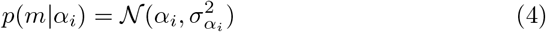

with a variance term, 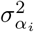 defined as a sine wave,

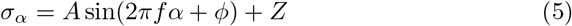

scaled to range between 0 and 1. The free parameters *A, f*, and *φ* to control the amplitude, frequency, and phase, respectively, while the value Z prevented the sine wave from reaching a value of 0. We selected a sine wave to operationalize the variance of the likelihoods for *α* as it is a simple model with few parameters that approximated observer changes in variance across *α* well.

The expected *α*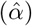 for any given trial was taken as the mean of the posterior distribution with variance 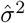. Model accuracy (percent correct) between the trial 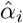 and the reference 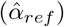 was defined as the probability returned by a cumulative normal distribution [45],

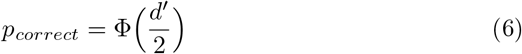

wherex

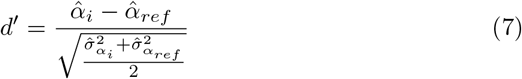

Discrimination thresholds were the Δ*α* value (difference between the test *α* and the reference a) that elicited 70.71% correct performance, as in our behavioral task. Model parameters (*A, f*, and *phi*) were optimized for each prior condition with *fminsearch* in MATLAB, minimizing the sums of square error between the model output and observer *α* discrimination thresholds, prior to and following adaptation.

Of the prior variants explored here, the best performing model had prior probability distributions that matched the adapting environment of observers (**Figure 4A**). The other model variants captured observer performance for their respective conditions well but were unable fit thresholds for other experimental conditions (see table 2 for the model *RMS_errors_* and Appendix A for the resulting model fits). Given these results, it appears that adaptation in modified environments, where the distribution of *α* has shifted towards steeper or shallower *α* can exert a change in the observer’s prior for *α* and reduce discrimination thresholds (**Figure 4A**). As would be expected from an adaptation paradigm, we also observe a systematic change in the likelihood variance across adaptation conditions (**Figure 4B**). To simplify visualization, **Figure 5** draws the envrionmental prior model standard deviation of the likelihoods for each environments. At baseline, the standard deviation of the likelihood peaks at an *α* of 0.7. It decreases as *α* steepens, with a trough around *α* of 1.6. Adaptation to a shallow environment reduced the width of the likelihoods for shallow *α* values while seemingly leaving steeper *α* values unaffected. Adaptation to a steep environment reduced the width of the likelihoods for *α* values between 1.0 and 1.3, mirroring the standard deviation of the likelihoods in the shallow adaptation condition.

**Figure 4:**
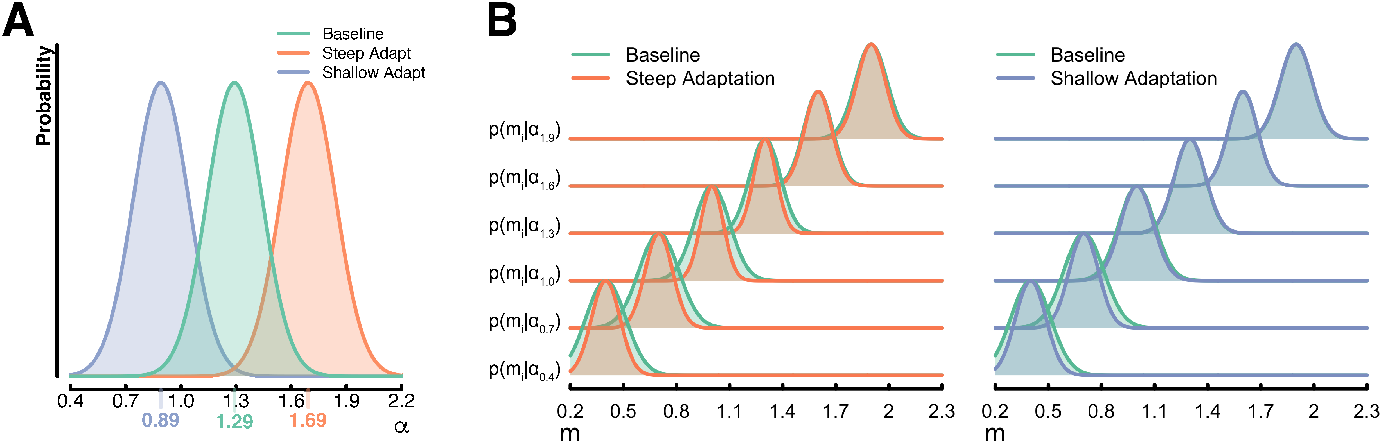
**A.** The environmental prior probability distributions of the best fitting model. **B.** Some of the resulting likelihoods from the model fitting procedure *p*(*m_i_* |*α_i_*) comparing the adaptation likelihoods to those of the baseline condition. We observed a narrowing of likelihoods for αs in the range of 0.4 to 1.3 following steep adaptation (with the narrowing being largest or most prominent for reference αs between 1.0 and 1.3). Only shallow αs (~ 0.4-0.7) showed a indication of narrowing following shallow adaptation.

**Figure 5:**
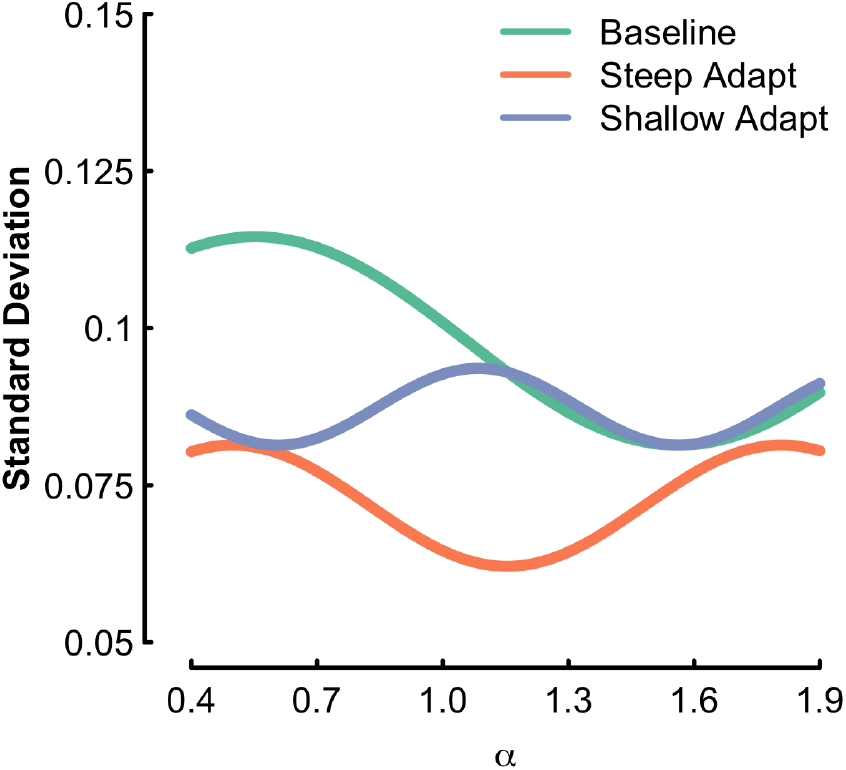
The standard deviation of the environmental priors model likelihoods across α. The standard deviation of the likelihoods was defined as a sine wave with amplitude, frequency and phase as free parameters.

**Table 2:** The R^2^, Root Mean Square error (RMS_*e*_) and the Akaike Information Criterion (AIC) calculated across all experimental conditions for each model investigated in this study. Only the environmental priors model and Steep prior had average deviations small enough to calculate R^2^ values for the model fits. Both models have similar AIC values, but the environmental priors model captured more of the variability in our data and returned a smaller average deviation than the steep prior model and is elected the best performing model.

| Model | $R^2$ | $RMS_e$ | AIC |
| --- | --- | --- | --- |
| Environmental Priors | 0.88 | 0.007 | -32.59 |
| Baseline Prior | - | 0.051 | -1.55 |
| Steep Prior | 0.59 | 0.014 | -32.62 |
| Shallow Prior | - | 0.065 | 0.43 |
| Natural Prior | - | 0.056 | -24.27 |

## Discussion

We measured discrimination thresholds to the slope of the amplitude spectrum of noise patches before and following immersion in a modified environment, all within an HMD. We found baseline *α* discrimination thresholds to have the same pattern across reference *α*s as is typically measured in more traditional laboratory equipment. Adaptation reduced *α* discrimination thresholds for reference *α* values near the average *α* of the modified environment. Specifically, immersion in a steepened world decreased thresholds for very steep reference *α* (i.e., 1.9), while immersion in the shallowed environment decreased thresholds for a reference *α* of 0.7. Behaviorally, our findings are similar to other visual paradigms (e.g., speed and color perception), which found improved sensitivity at values near the adapter [42, 43, 27]. These types of adaptation effects are thought to indicate an improvement in sensory encoding (an increase in the signal to noise ratio or a narrowing of the likelihood [42, 44, 43]. However, there is accumulating evidence that prolonged or repeated exposure to particular stimuli (i.e., adaptation) can have a meaningful effect on the prior probability distribution of observers, shifting them toward the values of the adapting stimuli [32, 31, 46, 47]. In order to determine whether an improvement in sensory encoding or a change in the expectation of the stimulus properties best explains our findings, we modeled our psychophysical results with a Bayesian observer similar to previous studies investigating the statistics of natural scenes [19, 31]. We built five model variants that differed in their prior probability distribution. We found the environmental prior model, with prior probability distributions representing the current distribution of *α*s, best-captured observer thresholds. Additionally, observer likelihoods were also affected by the adaptation paradigm. Shallow adaptation narrowed the width of the likelihoods near *α* values of 0.7, and a steep adaptation narrowed the likelihood for *α* values between 1.0 and 1.3. We find both a change in the likelihood and the prior explain the change in psychophysical responses. Our findings support that sensitivity to *α* is associated with the statistical structure or distribution of *α* in natural scenes and is sufficiently malleable to adapt in environments with novel *α* distributions.

Sensitivity to visual features is dependent on the probability of occurrence of these features in the environment of observers [32]. For example, the typical anisotropic sensitivity to orientation (i.e., the oblique effect) is not solely attributed to inhomogeneity in sensory noise for oblique orientations but also from their relatively lower occurrence in the environment compared to cardinal orientations [19]. Our modeling results suggest a similar phenomenon concerning *α* discrimination, which can—in part—be attributed to the occurrence of *α* values in natural environments. Following exposure to an environment with a modified *α* distribution increased sensitivity to *α* at reference values near the peak of the modified *α* distribution. To capture this effect in observer discrimination thresholds, the peak probability of the prior distributions had to match those of the distribution of *α*s in the environment. Changes in the prior probability distribution in the direction of the environment distributions have been reported for other visual paradigms [47, 31, 46, 32]. Exposure to higher speed stimuli over multiple days changed the speed prior of observers from one with high probabilities for low speeds (the low speed prior) to a prior probability distribution with a peak at higher speeds [47]. This change in the prior is also similar to effects shown by [31] where immersion in an isotropic world (in an HMD) flattened the prior for orientation [19].

However, changes to the prior alone do not appear to be sufficient to characterize the change in discrimination thresholds following adaptation fully, which was particularly evident in the shallow adaptation condition. The likelihood variance narrowed for *α* values near the mean *α* of the adapting environment, a reduction in measurement noise that is expected following visual adaptation. Previous Bayesian observer models have found the increased sensitivity following adaptation to stem from a narrowing of the likelihood distributions at the value of the adapter and not a change in the prior [42, 43]. It is plausible that the change in the likelihood following shallow adaptation contributed to reducing discrimination thresholds at a reference *α* of 0.7. Still, a change in the likelihood alone cannot adequately capture the discrimination thresholds across all three adaptation conditions. Model variants clamped to a single prior (i.e., baseline, steep, shallow, or the natural [5] priors – shown in Appendix A), while able to fit their respective experimental condition were unable to capture all experimental conditions.

In selecting a parameterization for the variance of our likelihoods, we opted for a sine wave function as it approximated the observer variance well in the current study and previous works [25]. The sine wave had the additional benefit of being defined by relatively few parameters (amplitude, frequency, and phase). A similar approach was used to determine the width of the likelihood when developing a Bayesian observer model for orientation content in natural scenes [19]. We had no explicit motivation in selecting this function to determine the variance of our likelihoods. A variety of simpler or more complex functions could also capture the change in likelihood variance across stimulus *α*. Our predominant interest in this study was to determine whether adaptation in modified environments could generate a shift in the prior of observers. Consequently, we did not explicitly design our experiment to accurately determine the shape of the function that defines measurement noise. Another paradigm better suited to determine measurement noise would likely aid in parameterizing the shape of said function across α. Given our current data, we opted to define the width of the likelihoods with a simple function that appeared to represent observer data well and sufficiently flexible to accommodate for changes in the likelihoods following either adaptation condition [42, 43].

Additionally, while we find evidence that a change in the likelihood contributes to the change in discrimination thresholds following adaptation, it is insufficient to characterize the change fully we observe following immersion in the modified environment. Our modified reality procedure subjects observers to the adaptation environment for an extended period (1 hour + 45 minutes of subsequent testing), which is significantly longer than traditional adaptation studies that adapt for periods of seconds or minutes. More extended adaptation periods are have been shown to target different visual mechanisms, which operate on longer-term visual representations than a short-term adaptation that alters mechanisms responsible for moment-to-moment fluctuations [48, 49]. Short adaptations periods are therefore unlikely to generate any meaningful change in the prior of observers. At the same time, a prolonged immersion in modified reality [31] or repeated exposure to certain stimuli over days [47] will stimulate the learning of novel stimulus properties / environmental statistics and effect a change in the prior probability distribution.

Finally, it is worth noting that psychophysical measurements of *α* discrimination thresholds are traditionally completed in laboratory environments with monitors (e.g., CRTs) that are large and of a sufficiently high resolution to display a wide range of spatial frequencies [24, 13, 25, 9]. Traditional CRTs appear not to be necessary when measuring sensitivity to the slope of the amplitude spectrum of natural scenes. While implementing psychophysical paradigms in HMDs is not novel [50, 31], the measurement of *α* discrimination threshold, which is dependent on the spatial frequency resolution of the display, is novel. We have demonstrated that discrimination thresholds to *α* can be measured accurately with the smaller screens of HMDs limited in the range of spatial frequencies they can display. The spatial frequency resolution of the Oculus Rift DK2 only reaches 5.5 cycles/° of visual angle. Still, *α* discrimination thresholds measured in the Oculus Rift DK2 showed the expected peak and trough for reference *α*s of 0.7 and 1-1.3, respectively. The peak and trough were also of similar magnitude to thresholds measured in other studies (using the same staircase methodology) [24, 25]. Human observers do not need a wide range of spatial frequencies to discriminate *α* and complete the task with relatively limited spatial frequency content.

## Conclusion

Discrimination thresholds to the slope of the amplitude spectrum (*α*) are subject to the recently viewed environment of observers. Spending a prolonged amount of time in a modified environment where the typical *α* values are steeper or shallower than the normal visual world of the observer will decrease discrimination thresholds to steep or shallow *α*, respectively. These effects were well-described by a Bayesian observer model, which showed a systematic shift in its prior probability distribution as a function of adaptation. The priors were shifted in the same direction as the adaptation condition. Our findings demonstrate that expectations on the occurrence of *α* values that result from a lifetime of exposure remain plastic, accommodating for the statistical structure of the recently viewed environment.

**Table 3:** Goodness of fit measures for each model explored in this study for all experimental conditions presented separately. Some models generated errors too large to calculate an *r*^2^, in these scenarios a – is used.

| Model | <u>Baseline</u> |  | <u>Steep Adaptation</u> |  | <u>Shallow Adaptation</u> |  |
| --- | --- | --- | --- | --- | --- | --- |
| | $r^2$ | $\text{RMS}_e$ | $r^2$ | $\text{RMS}_e$ | $r^2$ | $\text{RMS}_e$ |
| Environmental Priors | 0.966 | 0.004 | 0.669 | 0.012 | 0.911 | 0.004 |
| Baseline Prior | 0.966 | 0.004 | - | 0.087 | 0.817 | 0.005 |
| Steep Prior | 0.321 | 0.019 | 0.669 | 0.012 | 0.530 | 0.008 |
| Shallow Prior | - | 0.111 | 0. 729 | 0.011 | 0.911 | 0.004 |
| Natural Prior | 0.947 | 0.005 | - | 0.096 | 0.291 | 0.010 |

## Acknowledgements

We would like to thank Bruce C. Hansen for his insightful comments on this manuscript.

**Appendix A. Fits of the other models investigated in this study**

**Figure 6:**
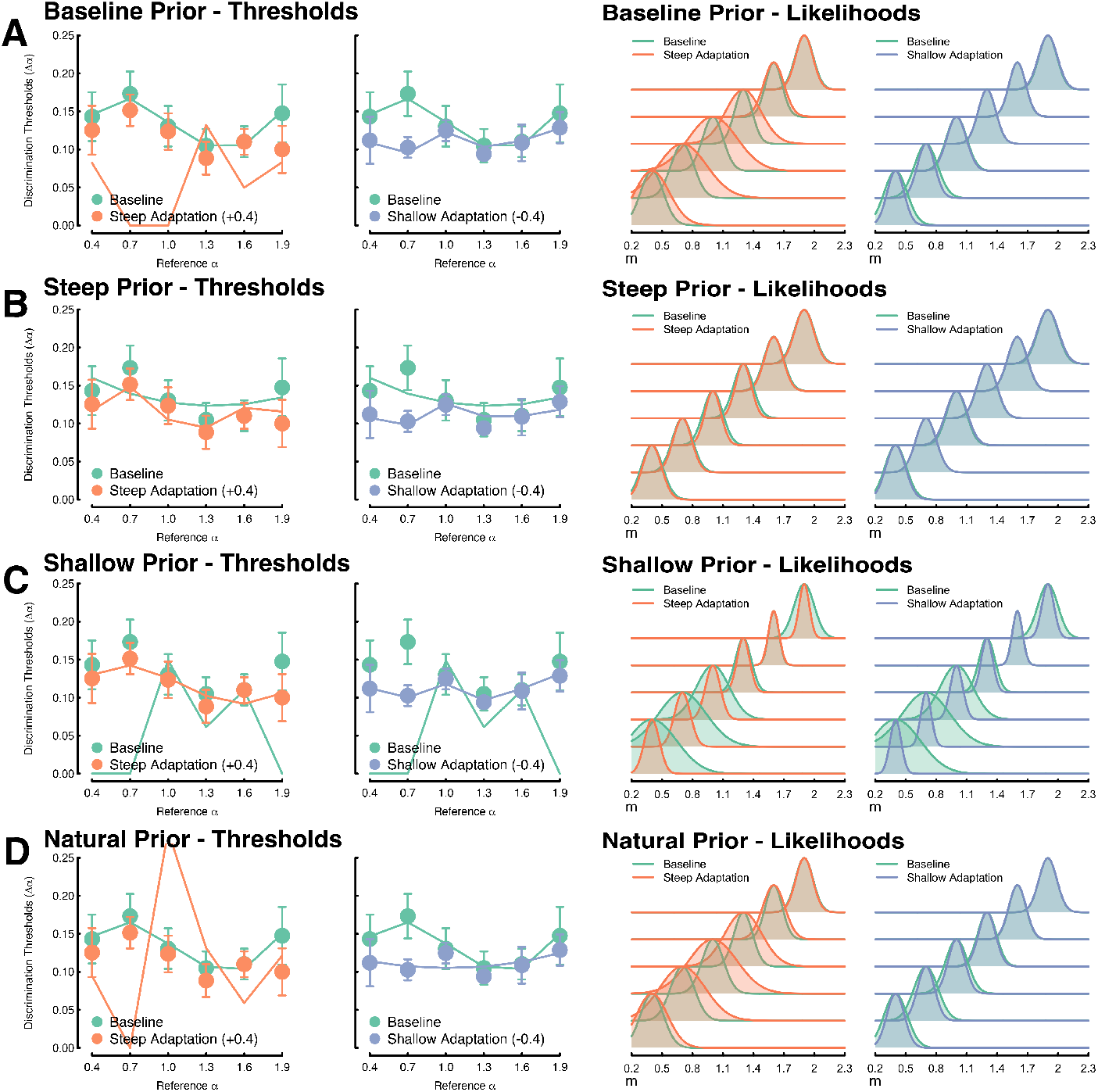
**A** The resulting fits and likelihoods of a Bayesian observer model with a prior probability distribution fixed to the baseline environmental distribution of *α* (*α* = 1.29). **B** The resulting fits and likelihoods of a Bayesian observer model with a prior probability distribution fixed to the steepened environmental distribution of *α* (*α* = 1.69). textbfC The resulting fits and likelihoods of a Bayesian observer model with a prior probability distribution fixed to the shallowed environmental distribution of *α* (*α* = 0.89). The resulting fits and likelihoods of a Bayesian observer model with a prior probability distribution fixed to the natural environmental distribution [5] of *α* (*α* = 1.08). Models A – C captured the discrimination thresholds of their respective experimental condition well, but failed when fitting other experimental conditions. The natural prior performed well with baseline discrimination thresholds but could not fit the other experimental conditions.

## Footnotes

1 Modifying the *α* of a natural image will alter its perceived focus: steepened images will appear blurred while shallowed images appear sharpened, which may aid in the discrimination task.

2 This low visual resolution is a consequence of having displays with a large field of views very close to the observers’ eyes, ~ 5cm.

